# Protein loop modeling and refinement using deep learning models

**DOI:** 10.1101/2021.11.03.467148

**Authors:** Feng Pan, Yuan Zhang, Chun-Chao Lo, Arunima Mandal, Xiuwen Liu, Jinfeng Zhang

## Abstract

Loops in proteins play essential roles in protein functions and interactions. The structural characterization of loops is challenging because of their conformational flexibility and relatively poor conservation in multiple sequence alignments. Many experimental and computational approaches have been carried out during the last few decades for loop modeling. Although the latest AlphaFold2 achieved remarkable performance in protein structure predictions, the accuracy of loop regions for many proteins still needs to be improved for downstream applications such as protein function prediction and structure based drug design. In this paper, we proposed two novel deep learning architectures for loop modeling: one uses a combined convolutional neural network (CNN)-recursive neural network (RNN) structure (DeepMUSICS) and the other is based on refinement of histograms using a 2D CNN architecture (DeepHisto). In each of the methods, two types of models, conformation sampling model and energy scoring model, were trained and applied in the loop folding process. Both methods achieved promising results and worth further investigations. Since multiple sequence alignments (MSA) were not used in our architecture, the energy scoring models have less bias from MSA. We believe the methods may serve as good complements for refining AlphaFold2 predicted structures.

## Introduction

Loops are protein fragments with an irregular structure connecting two secondary structural elements (usually α-helices and β-strands) and are important players in many biological processes. Their conformational plasticity plays a critical role in molecular recognition, allosteric control, binding of small molecules, and signaling [1-6]. As part of the overall protein scaffold, in some cases, loops adopt a relatively stable conformation in the folded state of the protein. Based on this static perspective, some efforts have been made to classify loop structures [7]. Conformational changes in loop regions, on the other hand, are frequently observed in various biological systems. For example, extracellular loops of G-Protein-Coupled Receptors (GPCRs) are critical for ligand recognition and binding [8]. Besides, the flexibility of loops is essential to drive enzyme function or regulation [9]. The concept of the energy landscape is used to better comprehend the dynamic nature of loops by providing a proper description of their conformational space [10]. Although a single loop can exist in numerous configurations, they are not all equally likely to be adopted. The likelihood of each feasible state is proportional to its associated free energy. However, due to its flexibility and irregularity, predicting the ground state for the loop has always been a challenge in the biological field.

Flexible loops are difficult to investigate and characterize structurally. As the most extensively used experimental approach for determining high-resolution protein structures, X-ray crystallography only offers static snapshots in certain experimental conditions[11]. Other experimental approaches, like X-ray solution scattering[12] or nuclear magnetic resonance (NMR)[13], can provide highly useful structural and kinetic information about these flexible regions, but only to a limited extent. Because obtaining a good atomistic description of the loop’s various conformations from experimental observations is extremely challenging, computational methods are an important alternative to the research.

Generally speaking, the process of loop modeling can be divided into three major stages: (1) conformational sampling, (2) energy scoring, and (3) post-processing or refinement. Over the last few decades, a variety of approaches to protein loop modeling have been proposed. Overall, there are three main classes: knowledge-based, *ab initio*, and hybrid approaches. Knowledge-based methods, also known as template-based, use structural repositories to extract observed loop conformations for a particular sequence and geometric information about the anchoring positions [14-17]. These approaches are computationally efficient in general since they do not depend on expensive simulations. However, they are constrained by the availability of acceptable loop conformations from known protein structures and only produce a limited number of solutions. *Ab initio* techniques, on the other hand, can sample a more significant proportion of the conformational space, for instance, by an exhaustive sampling of the loop’s torsional angles [18-19]. However, their computational costs are relatively high, and the methods are limited to short loops. For the hybrid approaches, many of them use small fragments from structural databases within an *ab initio* sampling technique [20-24] to achieve more balanced performance.

Most recently, deep learning methods using neural networks have dramatically impacted the computational biology field, including protein structure predictions [25-27]. AlphaFold2 was developed and achieved remarkable performance in CASP14[28], which is recognized to have solved the protein structure prediction problem at a similar level with experiments. Later RoseTTAFold was also developed, incorporating related ideas, and approach similar accuracy with AlphaFold2[29]. Even AlphaFold2 predicted the structures of many challenging protein targets near experimental resolution, but it was still weak for regions that have few intra-chain contacts, such as loops. Fig. 1 shows an example that AlphaFold2 performs prediction for the heavy chain on a Fab fragment (pdb id: 1mh5, chain: B). The overall prediction (red) had an excellent agreement with the native structure (blue), but the H3 loop region (yellow) had a space need to be improved. And the confidence score (plddt) of AlphaFold2 was also reduced at the H3 loop region compared with other parts. A recent DeepH3 based on deep learning was proposed, but it still reports limitations on long H3 loops with more than 12 residues [30].

**Figure 1:**
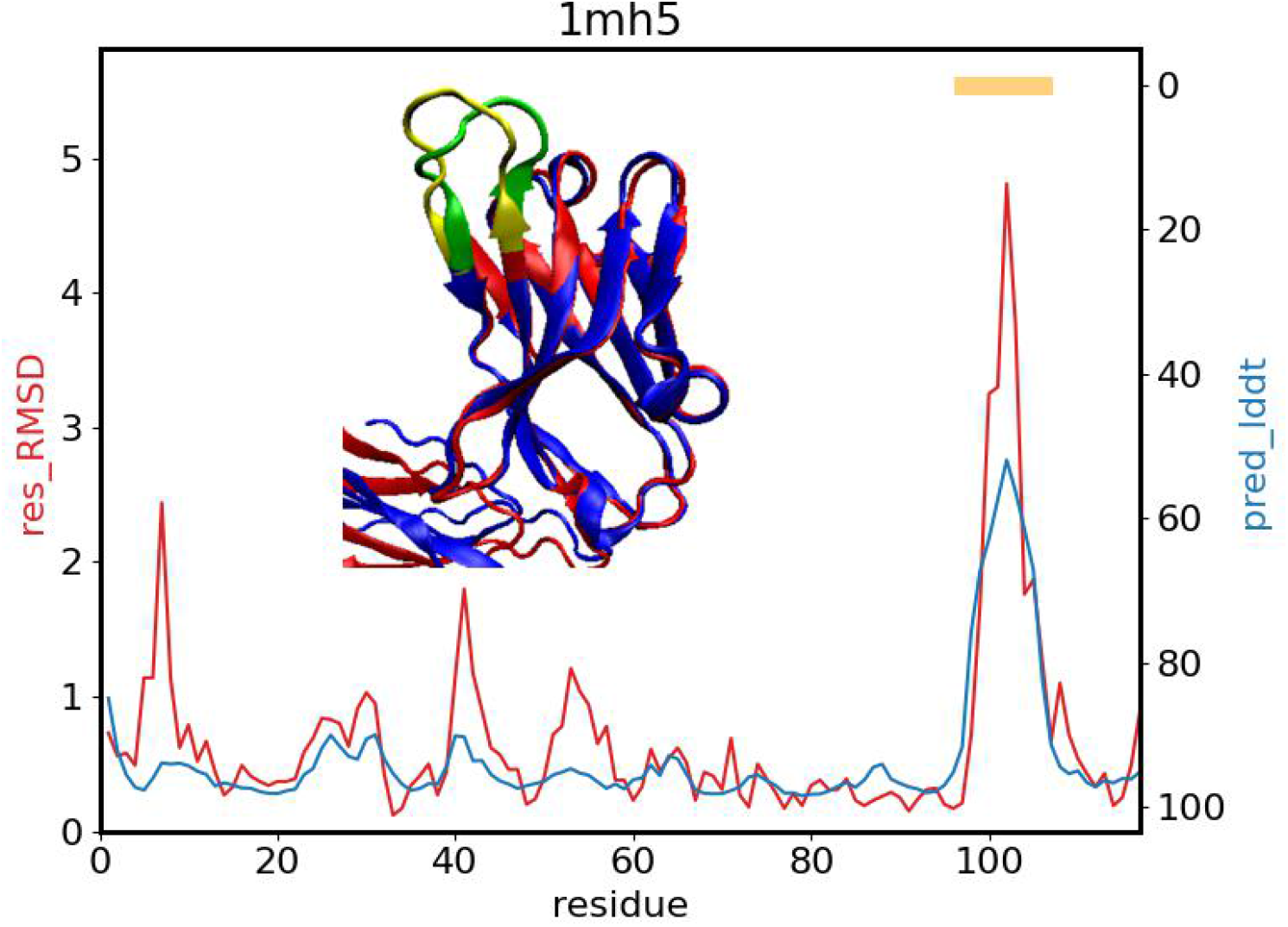
Snapshot of the predicted structure of 1mh5B (residue 1-117) by AlphaFold2 (red) aligned with the native structure (blue). The H3 loop region is colored green for native structure and yellow for modeled structure. The chart shows residue-wise RMSD (red) and predicted lddt (plddt from AlphaFold2, blue) as a function of residue index. The H3 loop region is labeled by an orange bar.

In this study, we designed and trained deep learning models for torsion sampling and energy scoring of loop modeling based on the 3D coordinate information as input. We used sequential chain-growth to perform simulated annealing folding given initial structure conformations. The test within training loops results in an average RMS of 1.39Å for loops with 16 residues.

On the other hand, because a change at a residue affects the inter-residue distances between all preceding and subsequent residues, using 3D coordinate information are highly interdependent.

The fragment needs to be predicted residue by residue without whole structure adjustment, leading to frustrate learning. Using 2D projections of 3D protein structure data, such as residue-residue contact map or distance map, is another way to represent the protein structure system. The advantage of such 2D projections includes translational and rotational invariance. Besides, the invariant information such as bone length and bone angle information which we relied on the machine learning model to learn from 3D structure, could also be implicitly included in the 2D projections. Plenty of previous work has been done to predict contact map from a multiple sequence alignment (MSA) and got decent structure prediction based on the predicted map, such as DeepMetaPSICOV[31], RaptorX-Distance[32], trRosetta[33], and AlphaFold[34] developed by DeepMind which significantly improved the accuracy of “free modeling” (no templates available) targets in CASP13[35]. Under the assumption that better contact information would lead to a better structure prediction, we propose a method to refine the atom-atom distance matrix from a decoy structure and do the structure refinement based on the refined distance map. Our distogram-guided refinement procedure improved 61.69% out of 154 fragments in our distogram refinement test dataset.

## Materials and Methods

### 1. Loop modeling

#### Cut-regrow sequential chain-growth

We use a sequential chain-growth method to generate the structure of the target loop when its starting and ending positions are decided. The detail of the procedure is shown in Fig. 2. With the starting N atom, the CA atom can be grown with given ω torsion. ω torsion is sampled based on its distribution in the database. Typically, except for proline(P), ω torsion should be around 180 degrees, while ω torsion of P could also be around zero. Therefore, we sample ω with Gaussian distribution around 180 degrees for all 20 amino acid types, but the special amino acid proline has some probability of being sampled around zero. Then C and CB atoms are grown with φ torsion sampled from trained neural network model and given torsion for CB. Similarly, the next N and O atoms are grown with ψ torsion sampled from the trained model and given torsion for O. When the growth comes to the CA atom within two residues of the N atom following the ending position, the analytical closure will be carried out to close the loop. It is not guaranteed that each growth can generate a closed loop, and we only keep those successful growths. For the bond lengths, bond angles, and torsion angles of CB and O, we use the value from DISGRO[21]. Those values are kept fixed except for analytical closure that we give small fluctuations to increase the success rate of growth.

**Figure 2.**
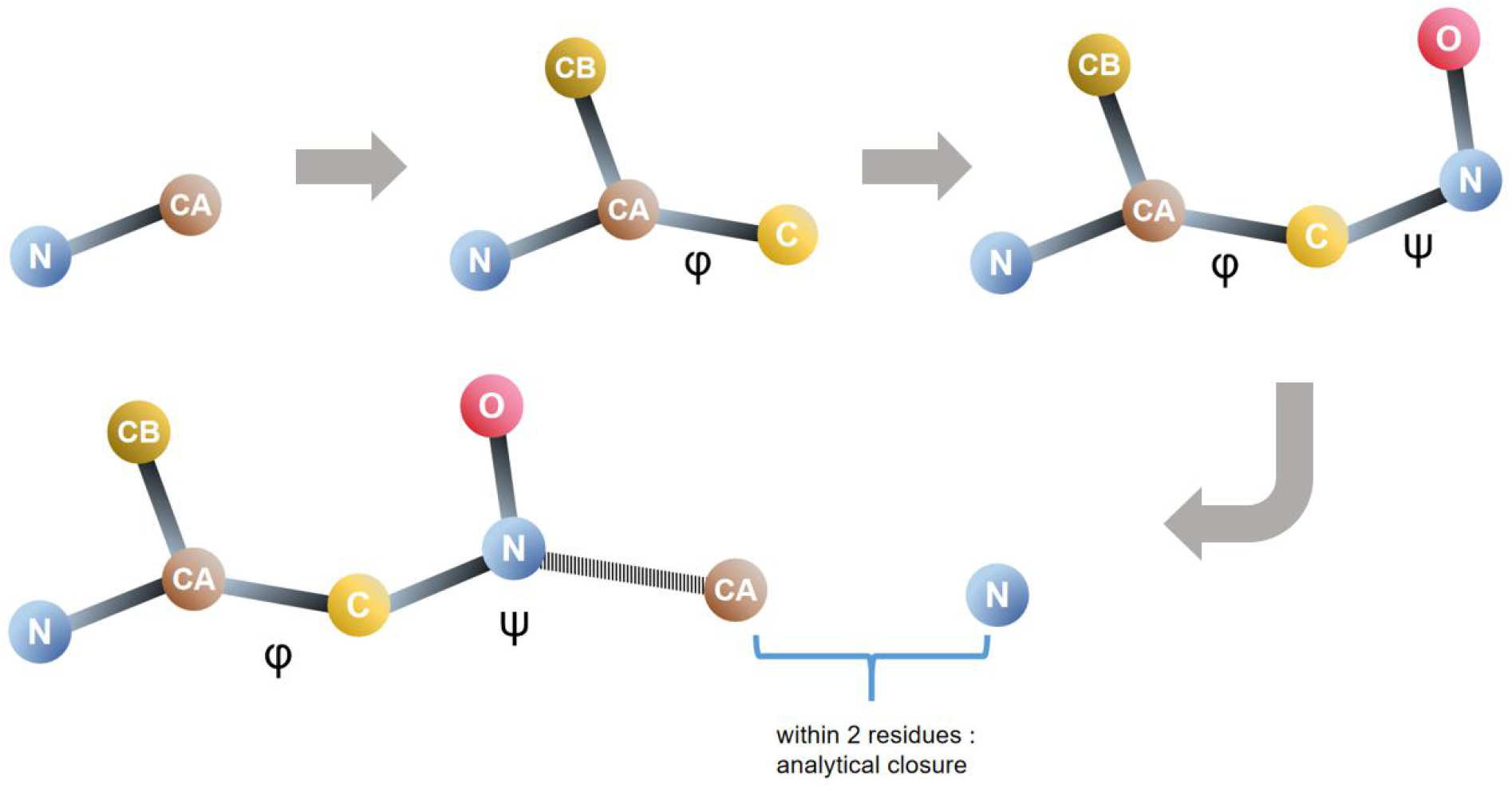
The flow of sequential chain-growth

The *Monte Carlo* cut-regrow scheme with simulated annealing is used to refine the whole fragment of the target loop. We randomly choose the cutting length and starting position to do regrowth many times following the sequential chain-growth process above. Then one regrowing structure with the lowest E is selected with the energy E as defined:

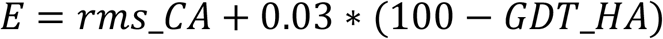

This E considers the importance of both rms and GDT in characterizing the structure, and the coefficient balances their weights. To accept or reject this regrowth, we use the Metropolis-Hastings probability regulated by simulated temperature:

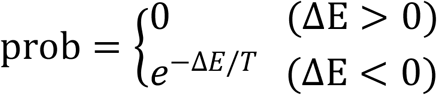

Where ΔE = E_f_ - E_i_, is the difference between the E after and before regrowth, and temperature T is decaying during the refinement process when a certain cycle of iterations is met as

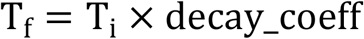

The parameters initial temperature T_0_ and decay_coeff need to be tuned to fulfill a fast and efficient refinement process.

#### Torsion model and Energy function model

##### Torsion sampling model

We designed a set of torsion prediction models (18 models in total) categorized by the length of growth (2 residues to 10 residues) and type of torsion angles (phi and psi). Take a length 10 fragment as an example, and the initial and final residues in this fragment were i and i+9. The input was two 3D gridded boxes, the center of one box (large box) located at the center point between two N atoms of the residues *i-1* and *i+10*, and the center of the other box (small box) located at CA_i-1_ for phi angle and N_i_ for psi angle. All the backbone atoms (C, CA, N, O) beyond the target fragment but within the box were included, which were labeled as one of 4 types. We also added a pseudo-CB atom to GLY. The CB atom at residue *i* had bond length for CA^i^-CB^i^ bond as 1.521 Å, bond angle N^i^-C^i^-CA^i^ as 110.4°, and dihedral angle N^i^-C^i^-CA^i^-CB^i^ as 122.55°. The CB atom beyond the target fragment but within the boxes were included as labeled as one of 20 types corresponding to 20 types of amino acid. The phi model also included the atoms N_i_ and CA_i_ in the target fragment, and this CA_i_ was labeled as the 25^th^ type to indicate the head of the grown fragment. The psi model used C_i-1_ as the indicator of the head, which was labeled as the 25^th^ type. The atom N_i+10_ was labeled as the 26^th^ type to indicate the tail of the grown fragment. The edge size of the small box was 16 Å for all models, and the size of large boxes were 18.0 Å, 18.0 Å, 20.0 Å, 24.0 Å, 28.0 Å, 32.0 Å, 34.0 Å, 38.0 Å and 40.0 Å for fragment length 2 to 10. With box size as BS, the input dimension was (N_b_, BS, BS, BS, 26). The box was gridded with each voxel of size (1 Å × 1 Å × 1 Å) for the small box and (2 Å × 2 Å × 2 Å) for the large box. We smoothed the input data using three-dimensional truncated Gaussian functions. Each heavy atom was represented by a Gaussian function, whose density was spread over the voxel the atom occupies and the 26 adjacent voxels. Each model took two boxes as input and output the predicted phi or psi for the first residue of the fragment.

The model consisted of two sets of sequential residual neural network (ResNet) blocks. Each set contains four ResNet blocks. Each block includes two layers of three-dimensional convolutional neural networks (CNN) with a kernel size of 3. The input boxes were fed into two sets separately and concatenated together after global average pooling. The vector was fed to a dense layer with 400 nodes, followed by the output layer outputs the predicted angle into 360 bins with the SoftMax as the activation function. We use the rectified linear unit (ReLU) as the activation function for all layers except the output layer. We use the categorical cross-entropy as the loss function and Adam for optimization. The learning rate is 0.0002, and the batch size is 100.

We choose 2187 protein structures with the sequence’s identity lower than 30% from PDB[36]. All the structures are determined by X-ray crystallography with a resolution better than 2.0 Å and do not have any DNA/RNA/UNK molecules. We use 80% of this dataset to train the torsion angle prediction model, 10% for validation, and 10% for testing.

The trained torsion models result in good efficiency in sampling the loops. We will use those models for all the simulations we did in this paper.

##### Energy scoring model

The input was the 3D conformation of the growth fragment captured by a series of 3D gridded boxes with an edge length of 18 Å centered at the CA atom of each residue. For each residue, the box was rotated to make its CA-C bond lying along the ‘ -x’ axis and its CA-N bond lying on the x-y plane. Each atom type is represented by a different channel (analogous to RGB color channels in images). Twenty-four channels are used, corresponding to atoms C, CA, N, O, OXT, and 20 types of CB atoms. All the boxes were fed into four ResNet blocks, and each block consists of two CNN layers with a kernel size of 3. The output vectors were fed into a bidirectional long short-term memory (LSTM) network, followed by global average pooling, two dense layers with node numbers as 128 and 32, and then an output layer with a linear activation function to predict the RMSD score. We use ReLU as the activation function for all layers except the output layer. The learning rate is 0.00005, and the batch size is 50.

##### *Monte Carlo* beam search

When generating near-native structures and structures based on the target distogram mentioned below, we use the *Monte Carlo* beam search algorithm and sequential chain-growth, as illustrated in Fig. 3.

**Figure 3.**
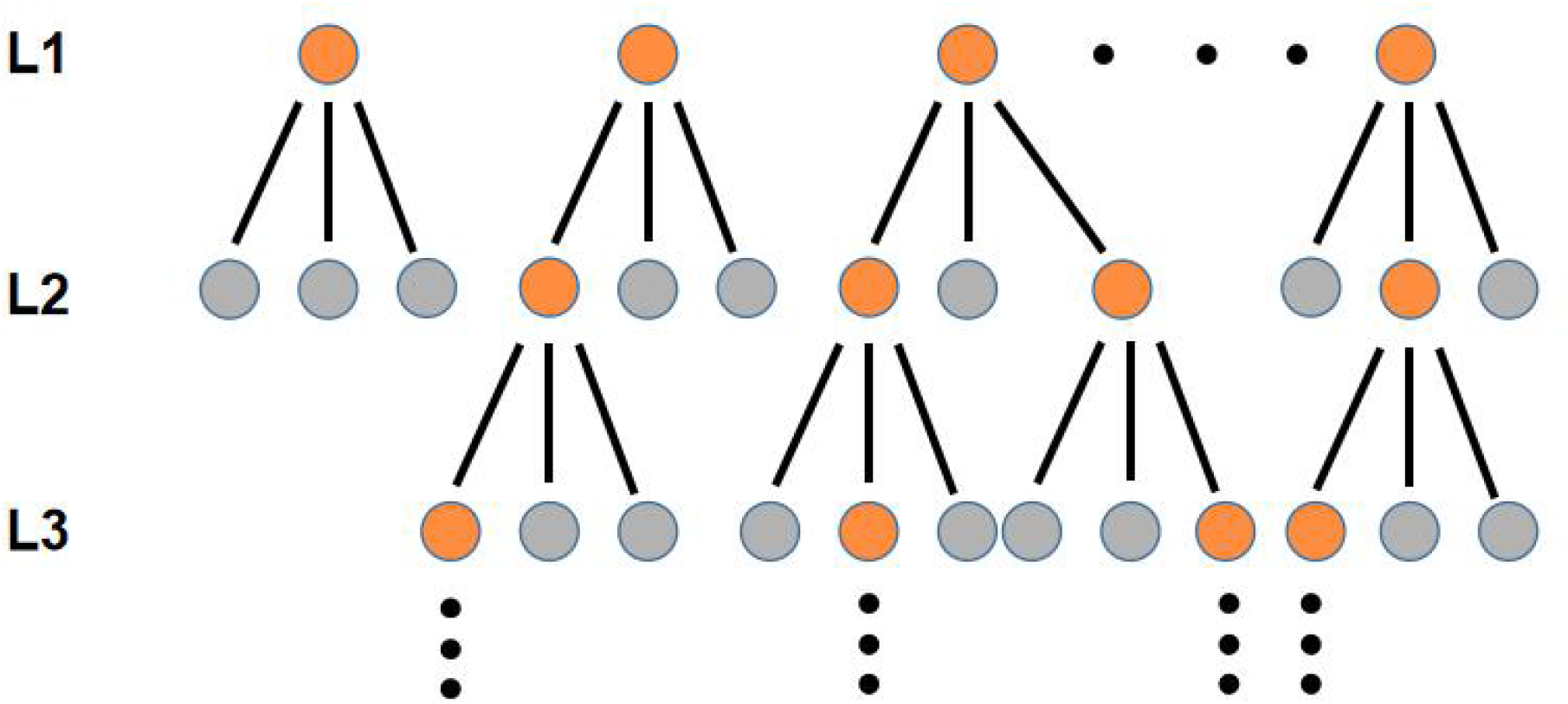
*Monte Carlo* beam search method. In this figure N_s_ = 3 and N_k_ = 4.

For the algorithm, we need a parameter ξ to evaluate each level and two parameters to control the sampling: number of samples N_s_ and number to keep N_k_. Typical for chain-growth, we can choose evenly distributed torsion angles between 0 and 360 degrees in level 1 and choose the best N_k_ nodes that minimize ξ. In level 2, we use the N_k_ nodes to continue N_s_ growth for each, leading to total N_s_*N_k_ samples. Then we choose the best N_k_ nodes out of N_s_*N_k_ samples that minimize ξ and move on to level 3. It is noticed that some nodes from level 1 may be eliminated like the first node in Fig. 3, and some may generate multiple sub-nodes. Besides, to sample more efficiently, it is best to sample near the best torsion angle minimizing ξ within a range η. Here, the three parameters N_k_, N_s,_ and η control the quality of sampling. To get more diverse sampling and a higher chance to find the global minimum of ξ, larger N_k_, Ns, and η are better choices while it may be costly in computation. So, the parameters need to be tuned to have good performance in practice.

#### Training of energy model

In this work, we use two kinds of data to train our energy function model, which are named FOLD_N and FOLD_M.

##### FOLD_N

FOLD_N represents native structure-oriented folding data. To generate this kind of data, we use the RMS_CA and GDT_HA with respect to the native structure to calculate E during the cut-regrow sequential chain-growth process. And for each iteration of cut-regrow, if the new fragment is accepted, we record the 3D structure as one input data for our energy function model. Data of different cutting lengths will be mixed to train one model, which is more efficient than training different models for different cutting lengths. The initial structures are generated using DISGRO. Since this is native structure-oriented, most of the input data will lead to “good outputs”, which means low E states.

##### FOLD_M

The problem of only using FOLD_N data is obvious. It will primarily cover the near-native space but not the folding space using models. To solve this, we introduced FOLD_M that is similar to manual reinforcement. For FOLD_M, we use the model trained based on only FOLD_N data to do cut-regrow sequential chain-growth, calculate E using the model’s prediction, and get the input data similarly as in the FOLD_N part. So, the true folding space using the model will be included to enhance the model. Technically, this iteration can be repeated many times to improve the model, while we only do it once in this work considering computation cost.

##### Asymmetric loss function

In actual practical trials, the model tends to be underestimated easily when we use the MSE loss function. That means, for many states with high real E scores, the model will predict it with low E scores. To deal with this, a Lin-Lin asymmetric loss function is introduced as follows

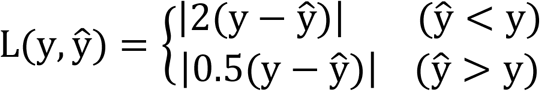

where ŷ is the predicted value, and y is the true value. By this definition, when the prediction is lower than the truth, the loss will be exaggerated while reduced *vice versa*.

### 2. Fragments of protein structure refinement guided by distogram

Instead of using a 3D structure as an input, we also design a method using an atom-atom pairwise distance matrix as an input to refine a fragment. We train a sampling model and two energy function models to refine the backbone structure of six-residues fragment pieces of protein structures described below.

#### Data preparation

To refine a length six fragment from a 3D protein structure, we first calculate the distance of the backbone atoms of this fragment with all the backbone atoms in the 3D structure and pick the nearest 100 residues spatially. We only consider the backbone atoms C, CA, N, O, and CB, where a pseudo-CB atom is added to GLY or any residue missing CB. We calculate the distance matrix of these 500 atoms with the 30 atoms in the target fragment and then convert this distance matrix to one hot (500, 30, 81). We divide 0-10 Å into 50 bins, 10-25 Å into 30 bins, and all the distances larger than 25 Å as one bin. We include the atom type for C, CA, N, O, and 20 types of CB information for each atom pair and convert it to one hot (500, 30, 48). We also calculate the absolute residue index distance for each atom pair (*i, j*) and calculate a residue distance score Rd_score_ (500, 30, 1), which is defined as:

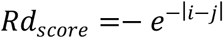

In this setup, we have input dimensions as (500, 30, 130), and the sampling model will output a distance histogram (distogram) in dimension (500, 30, 81). We will use this predicted distogram to generate a refined fragment.

We choose 2187 protein structures with the sequence’s identity lower than 30% from PDB [36]. All the structures are determined by X-ray crystallography with a resolution better than 2.0 Å and do not have any DNA/RNA/UNK molecules. We then split each protein chain into six-residues pieces and get 81231 fragments in total. We use the torsion sampling model and the *Monte Carlo* cut-regrow scheme to generate 25 sampled structures for each fragment. Another 20 near-native structures are generated using *Monte Carlo* beam search with real RMS as ξ, and only those with RMSD larger than 0.6 Å are kept. We randomly pick 80% protein chains (1749 protein chains, 129993 sampled fragments) as training data, 10% (218 protein chains, 15817 sampled fragments) as validation data, and the rest 10% (220 protein chains, 16620 sampled fragments) as test data.

#### Sampling model

Figure 10(a) shows that the sampling model (MS) is 12 layers convolutional neural network. The input has dimensions (N_b_, 500, 30, 130), where N_b_ is the batch size, and 130 is the channel number. In the first layer, we apply a 1 × 1 filter, which will create a linear projection of the features, decreasing the feature number to 64. In all the convolution or transpose layers of this model, we set stride as one and proper padding size to make the output dimension (H, W) consistent with the input. The result will be fed into two convolution layers with filter size 2×2 and channel number increasing from 64 to 128 and 256. The result will be fed into three convolution layers with filter size 3 × 3, channel numbers 256, 128, and 64. Six transpose convolution layers are followed with the symmetric setting with the convolution layers, as shown in Figure 10 (a). The output finally will be fed into a 3×3 convolution layer with channel number 81, which will generate a refined distogram of the fragment with dimension (N_b_, 500, 30, 81). The neural network is constructed using the Keras library [37]. We use the rectified linear unit (ReLU) as the activation function for all layers except the output layer, where we use the SoftMax activation function. We use the categorical cross-entropy as the loss function and Adam for optimization. The learning rate is 0.002, and the batch size is 20.

Our model is trained to generate a better distogram ŷ from a distance matrix ***x*** of a low-quality structure:

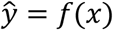

To test the performance, we first calculate the predicted distance matrix 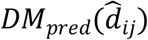 with dimension (500, 30, 1) from the output distogram ŷ with dimension (500, 30, 81). Then we calculate the dRMSD of 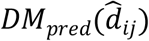 with the distance matrix of the native structure *DM*_*nat*_(*d*_*ij*_). All the distances larger than 25 Å will be converted to 25 Å before comparison.

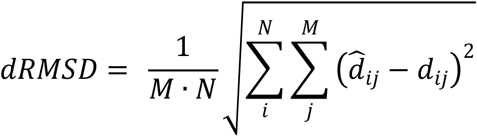

#### Energy model

As shown in Figure 10 (b, c), we have trained two models to predict the root mean square distance (RMSD) of a protein structure fragment compared with native structure by giving distance matrix, atom types, and residues separation. The input of energy models is the same as the sampling model, and the output is the predicted RMSD.

The baseline model (ME_resNet) contains three convolutional residual network (resNet)[38] blocks, which consists of three 2D convolution layers followed by a batch normalization layer. The size of filters in each residual block is kept the same but increases between blocks, which are 1×1, 2×2, and 3×3. The channel number is 256 for all convolution layers. We use global average pooling to convert the 2D layers to 1D vectors and then feed the vector to a dense layer with 400 nodes. We use ReLU as the activation function for all layers except the output layer, where we use the linear activation function to output the predicted RMSD. The learning rate is 0.002, and the batch size is 20.

The other energy model (ME_resNet_SD) is trained by using a dilated convolutional residual network with stochastic depth [39] shown in Figure 10 (c). The “survival” probabilities of the six blocks are set as 0.92, 0.83, 0.75, 0.67, 0.58, and 0.50. Each dilated residual block contains a 2D CNN layer with batch normalization, then follows a 2 × 2 average pooling layer to reduce the dimension of the matrix from (N_b_, 500, 30, 128) to (N_b_, 250, 15, 128). The size-reduced matrix will be fed into a 3×3 2D dilated CNN layer with batch normalization. And then, a 2D transpose CNN layer will be used to resume the matrix size back to (N_b_, 500, 30, 128). The dilated rates for each block are 1, 2, 4, 1, 2, 4, respectively. We use global average pooling to convert the 2D layers to 1D vectors and then feed the vector to a dense layer with 400 nodes. We use ReLU as the activation function for all layers except the output layer, where we use the linear activation function to output the predicted RMSD. The learning rate is 0.002, and the batch size is 20.

#### Refinement pipeline

Figure 11 shows the pipeline that we use the sampling model (MS) and energy model (ME) to refine a low-quality input fragment. The input is the 3D protein structure and the target fragment residue numbers. The refinement will be repeated N iterations, and each iteration is shown as in a simulation unit (SU). In each SU, the distance matrix and input feature will be calculated and fed into MS to get the refined distogram. Based on this refined distogram, we generate ten sampled 3D structures of the fragment using the *Monte Carlo* beam search with distance to target distogram as ξ. Then we will use ME to pick the one with the lowest predicted RMSD and output the structure. We repeat the SU 100 times or terminate the process once no structure is generated based on the predicted distogram. We randomly pick 154 fragment conformations (from 99 pdb structures). We make two experiments: one with MS_EP34 (ME model, epoch 34) to generate refined distogram, and ME_resNet_EP21 (ME_resNet model, epoch 21) to control output quality for each step; the other one with MS_EP34 for sampling and ME_resNet_SD_EP13 (ME_resNet_SD model, epoch 13) for quality control.

## Results and Discussion

### Loop modeling using chain-growth

#### MSE loss function VS asymmetric loss function

Fig. 4 shows the scatter plots of predicted E score against real E score of test data set for models of 10 L16 (length 16) loops using either MSE or asymmetric loss function. It shows that both models give good linear fitting with R coefficients 0.902 and 0.913. However, the slope for the model using the MSE loss function is 0.515, which means most real E scores are underestimated in prediction. For the model using asymmetric loss function, the slope is 0.999, that is very close to one. There are a few data points that are overestimated on the large real E score side. But since the number of these data points is small and overestimation is not as misleading as underestimation in refine regrowth, this overestimation can be ignored.

**Figure 4.**
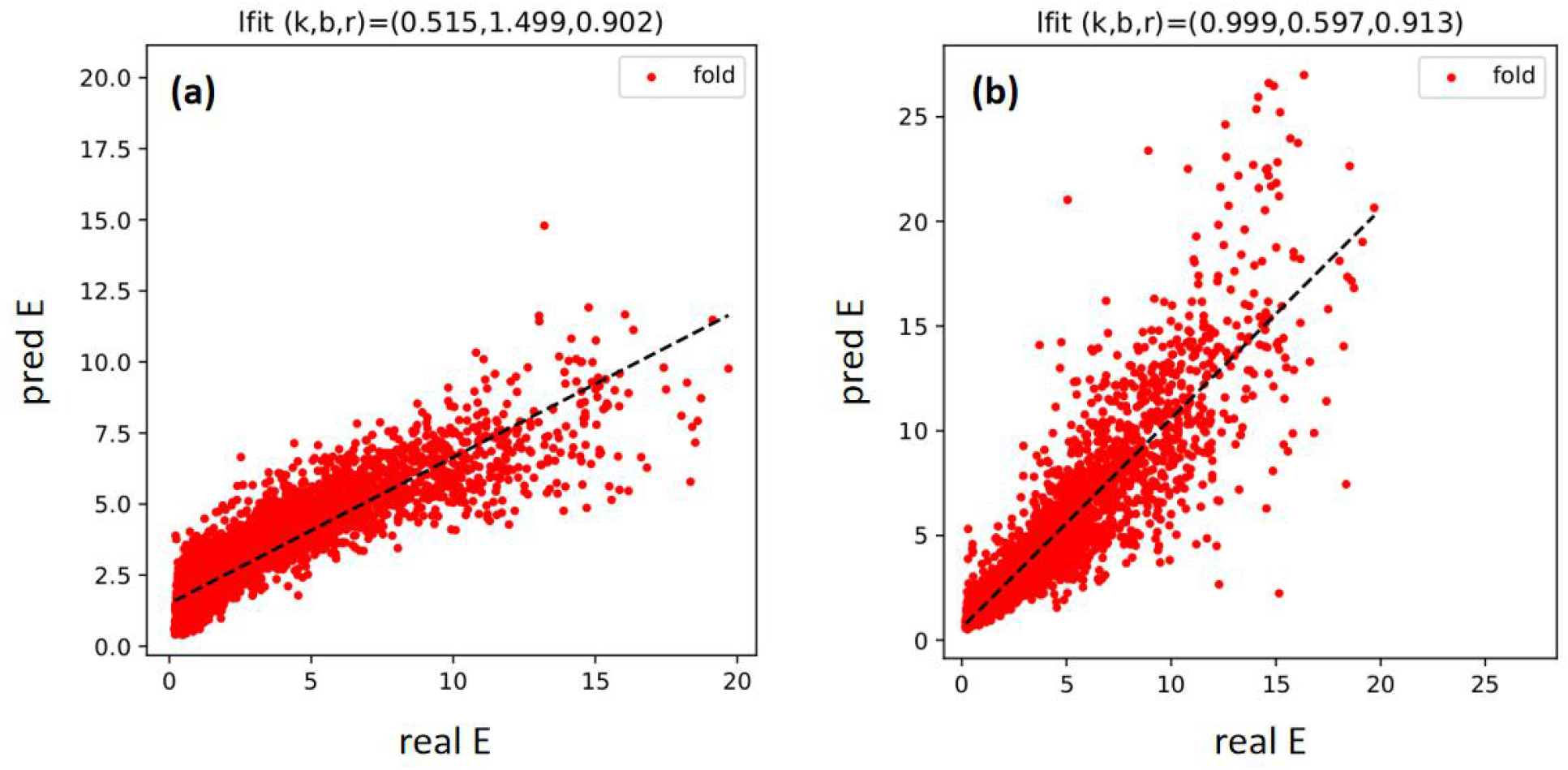
Comparison of results between MSE(a) and asymmetric loss function(b), the linear regression results are labeled on top.

#### Model of 42 L10 (length 10) loops with FOLD_N data

As stated in the methods section, a model of 42 L10 loops was trained based on the model’s good performance of 10 L10 loops. Only FOLD_N data was used here, and an asymmetric loss function was used. The cutting length was chosen from length 3 to 6. The training was finished up to 100 epochs, and Fig. 5 shows the training and validation loss against epochs. It is shown that the training loss keeps decreasing, and validation loss decreases with fluctuations. It is found that epoch 96 gives the lowest validation loss with 0.102. We plot the predicted E against the real E score in Fig. 6, and it shows good linear fitting between them for both the training set and test set. Then 10 loops were selected to do a cut-regrow folding test up to 1000 iterations. The initial structures were not chosen with the ones used in training. We set the initial temperature to 1 and the decay coefficient to 0.9. As seen in Fig. 6, the real RMS of all the 10 fragments drop very fast in the first 200 iterations and still get small refinement until the end. The average RMS of 10 fragments at the end of refinement is 0.673Å, which is a very low number for length 10 loops

**Figure 5.**
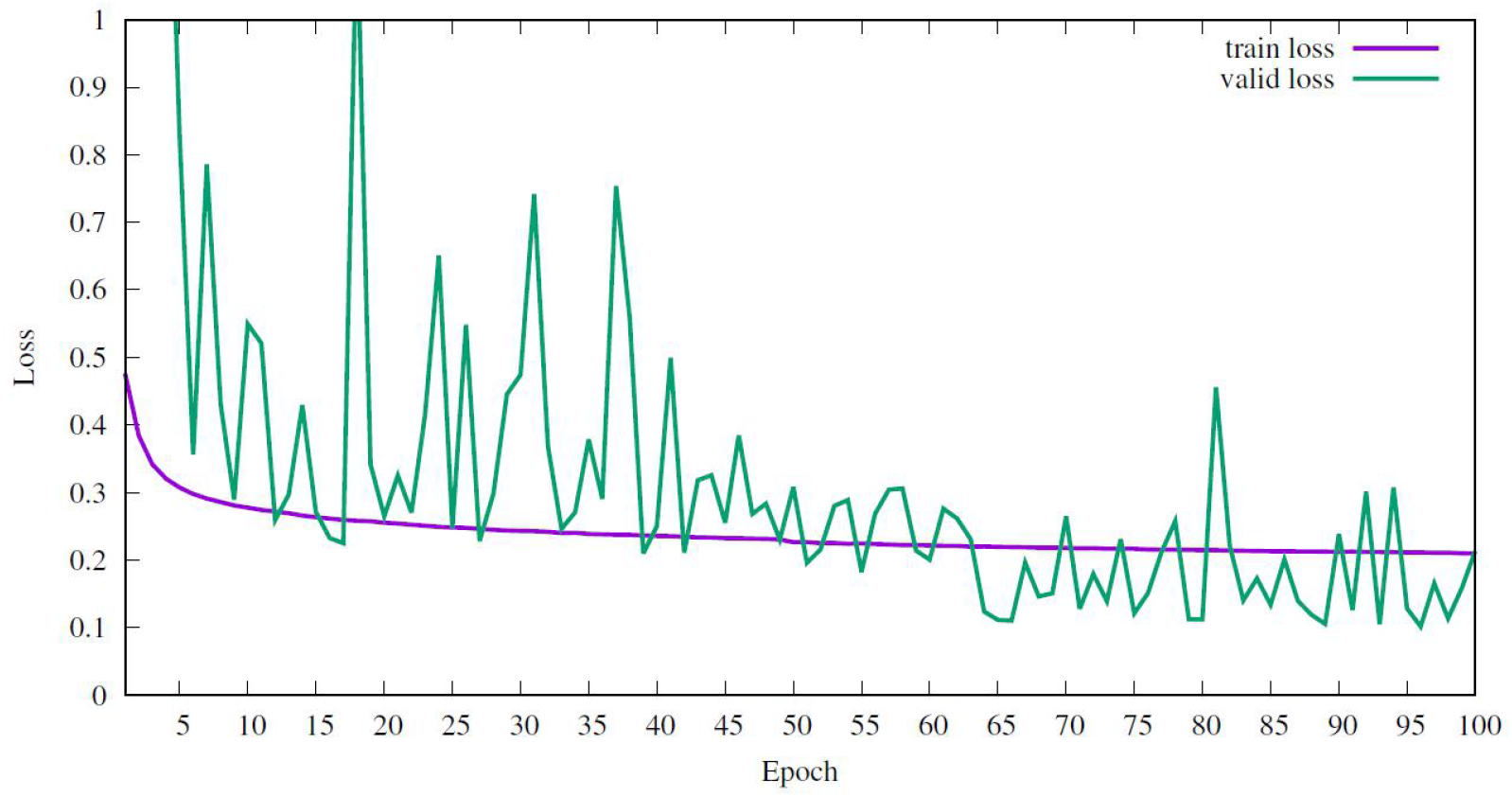
The training and validation loss for a model of 42 L10 loops

**Figure 6.**
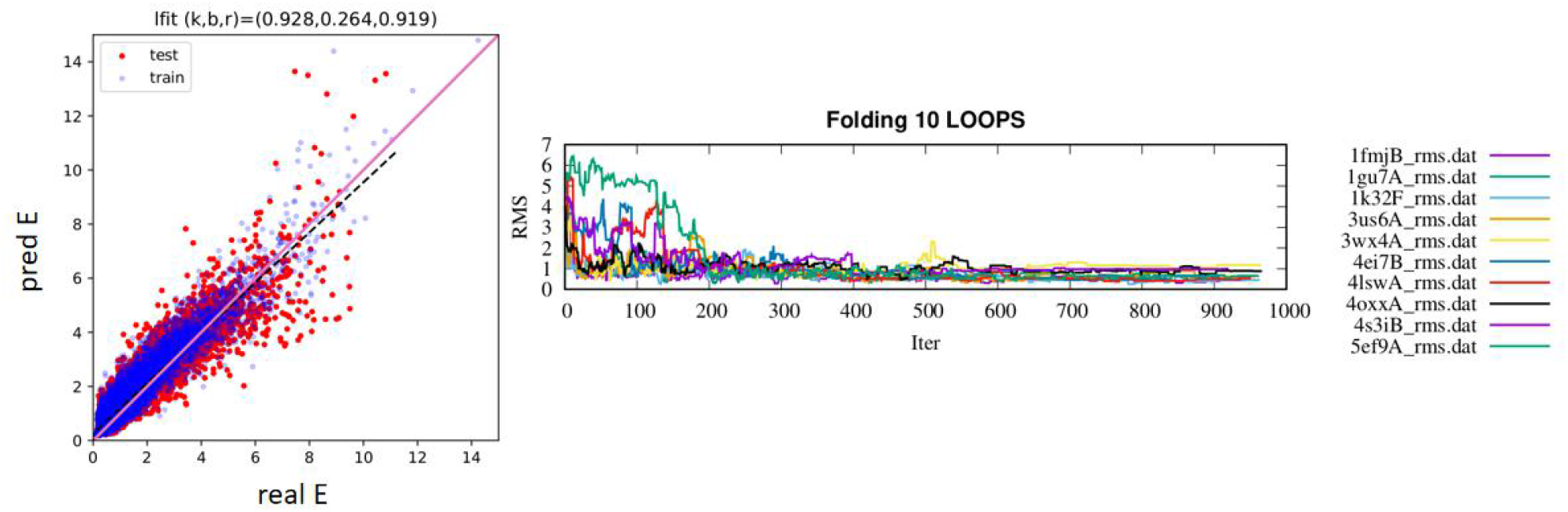
The E score prediction(left) and simulated folding results(right) of the model of 42 L10 loops

#### Model of 1000 L16 (length 16) loops

The model of 42 L10 loops shows good performance, but it works only within the loops selected in the training set. To train a more generalized model, we need to include more loops for our training data set. Therefore, the model of 1000 L16 loops was trained using both FOLD_N and FOLD_M data. We split 1000 loops into three datasets. For the first dataset D1, 800 loops are included with around 1600 FOLD_N data for each. For the second dataset D2, 100 loops are included with around 9000 FOLD_N data for each. For the third dataset D3, 100 loops are included with around 1600 FOLD_N data and 4500 FOLD_M data for each. FOLD_M data is generated based on a pre-trained model using only FOLD_N data. The total number of data is around 2.74M. The cutting length is from 6 to 10, asymmetric loss function was used, and the training finished up to 33 epochs. The training and validation losses are shown in Fig. 7. Like the model of L10 loops, both the training and validation losses are decreasing with some fluctuations for validation loss. The minimum validation loss is found at epoch 32 with 1.7535.

**Figure 7.**
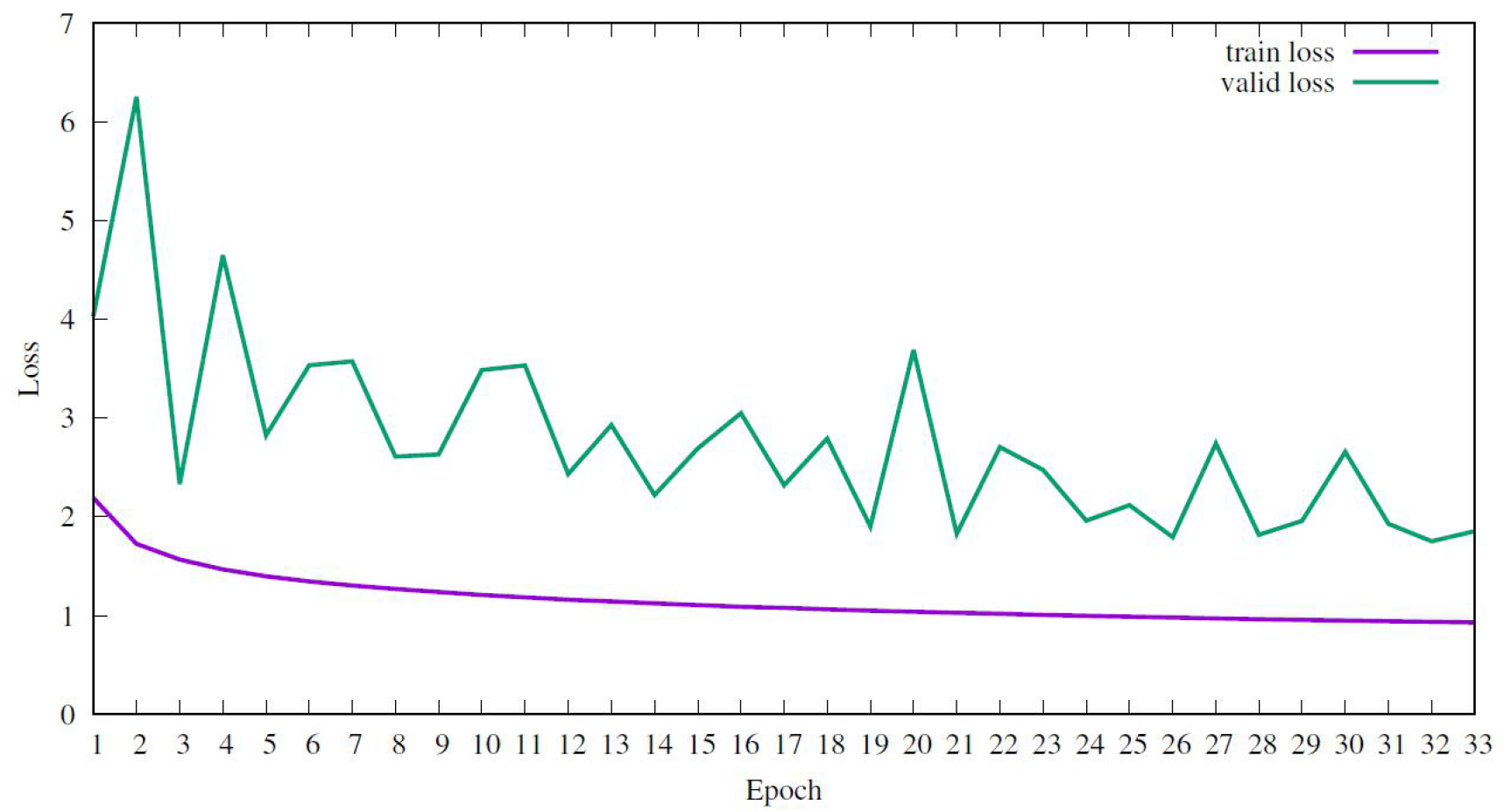
The training and validation loss for the model of 1000 L16 loops

Fig. 8 shows the predicted versus real E scores for the training set and test set and all three datasets. All three datasets give a good linear correlation for training and test sets. D2 has a better correlation than D1 because more FOLD_N data are used in training. The inclusion of FOLD_M data significantly improves the prediction accuracy, as D3 gives the largest R coefficient even though the total amount of data for D3 is less than D2. For the test set, the fitting coefficient is slightly smaller than the training set.

**Figure 8.**
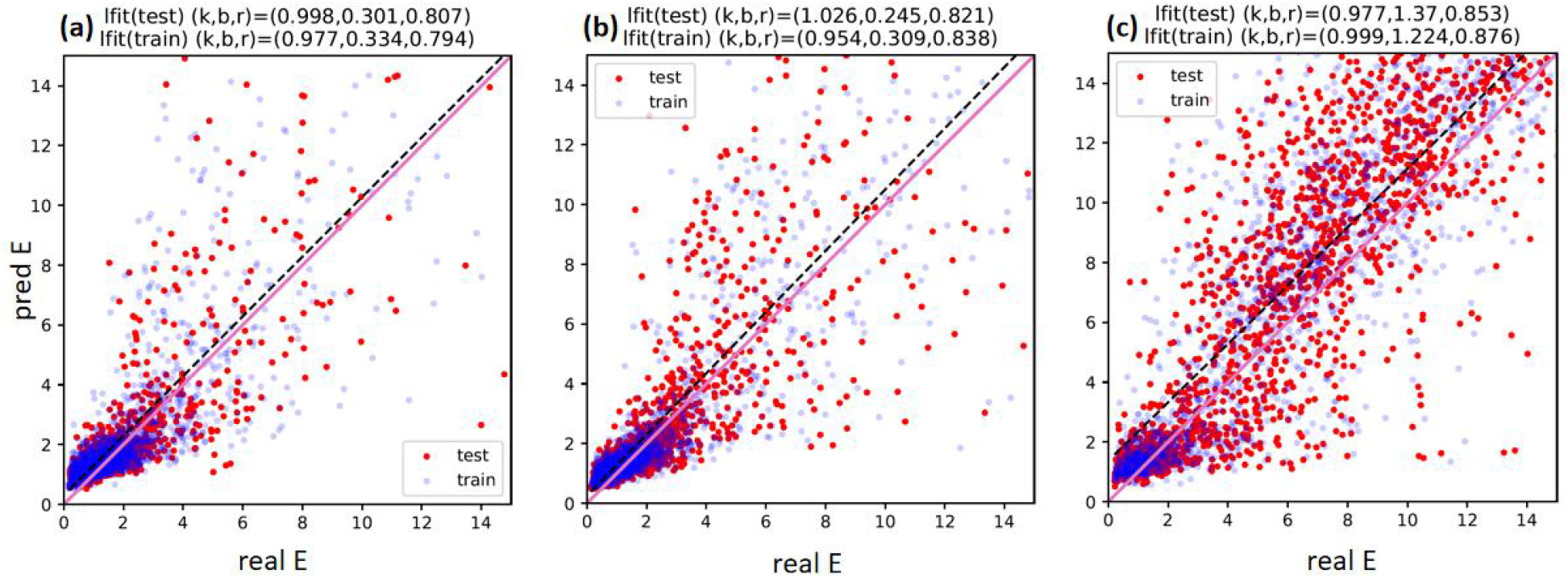
Linear fitting between predicted and real E scores for D1(a), D2(b), and D3(c) dataset.

To do the cut-regrow refinement, we picked ten loops from each dataset of D1, D2, D3 together with another dataset, D4, where ten loops not included in the training are selected. The initial temperature was set to 2 and the decay coefficient to 0.8. The real RMS against iterations are shown in Fig. 9. For D1, D2, and D3, most loops can be refined to a smaller RMS. At the end of refinement, the average RMS turns to be 3.21Å, 2.41Å, and 1.39Å for D1, D2, and D3. More FOLD_N data and including FOLD_M data can both help to improve the refinement process. With the use of FOLD_M data, the loops can be refined with an average RMS 1.39Å, which is a small number for L16 loops. However, for D4 loops, the results are not as good, with average RMS of 6.60 Å. This indicates the model is still not generalized very well. However, compared with the average initial RMS of 9.08 Å, the average RMS still can be improved by 2.48 Å.

**Figure 9.**
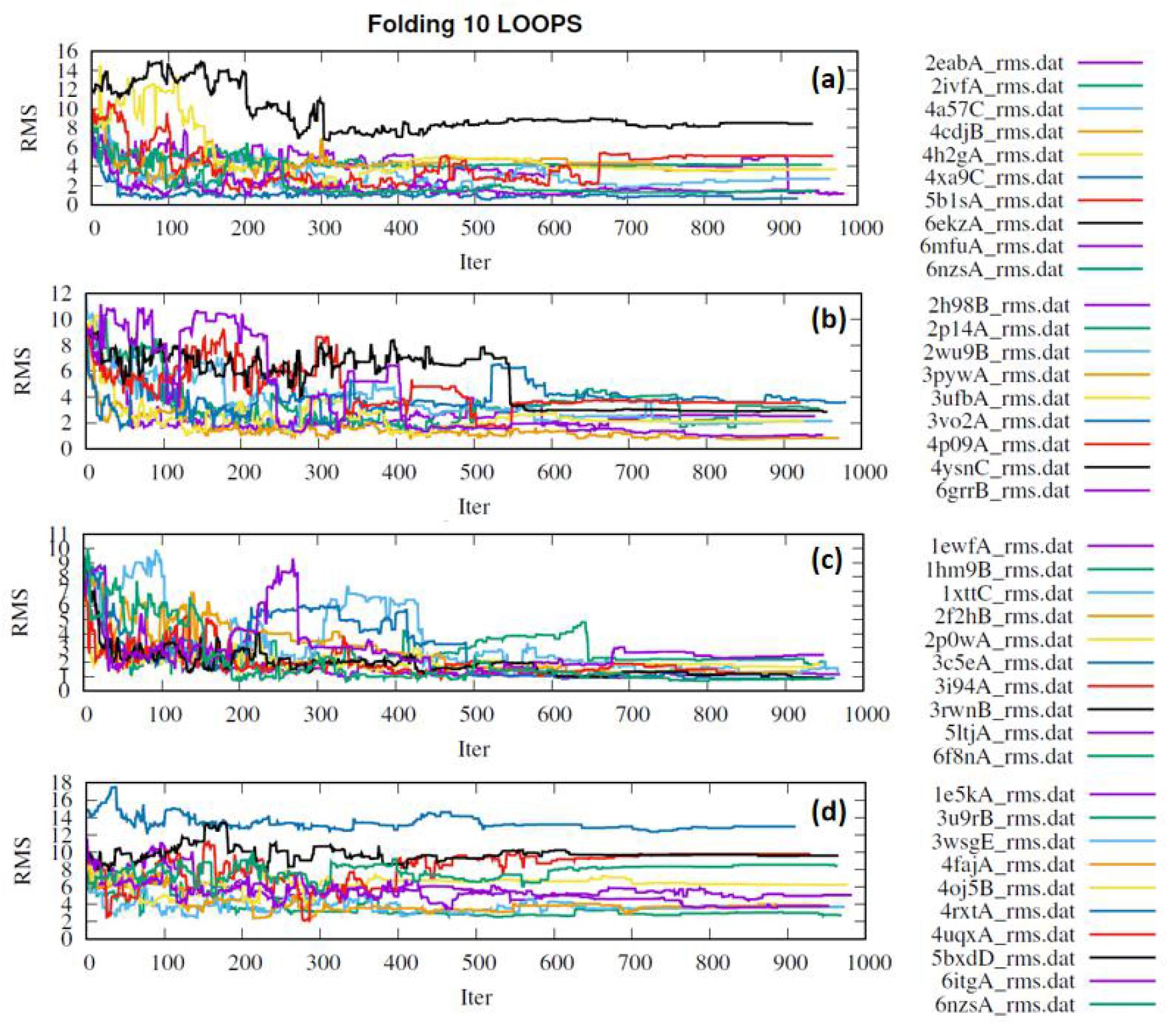
The simulated annealing folding curves up to 1000 iterations for selected loops from D1(a), D2(b), D3(c), and D4(d) dataset

**Figure 10:**
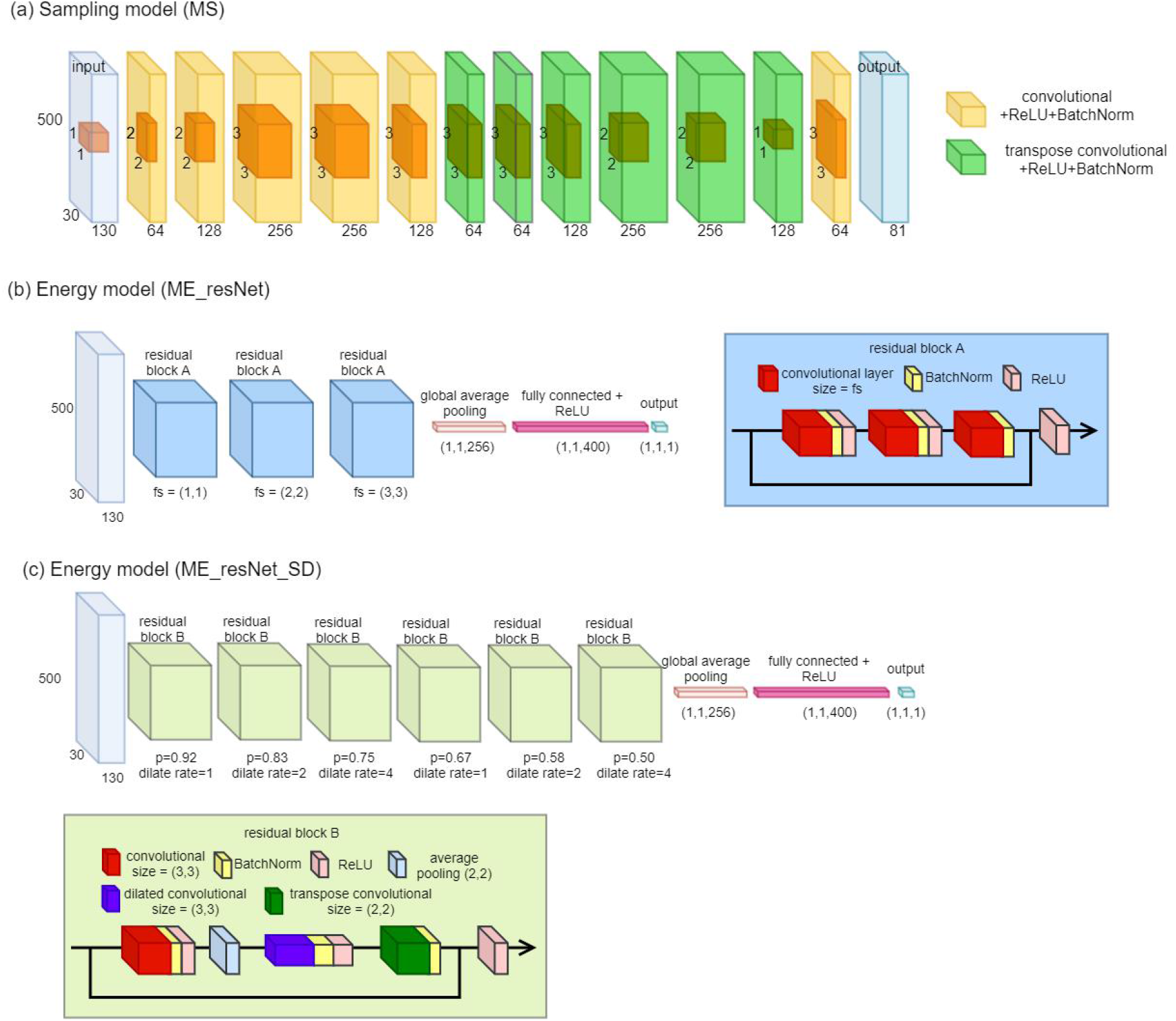
Architecture details. (a) Sampling model. (b) Energy model ME_resNet and the residual block A shown in the right box. (c) Energy model ME_resNet_SD and the corresponding residual block B are shown in the bottom box.

**Figure 11:**
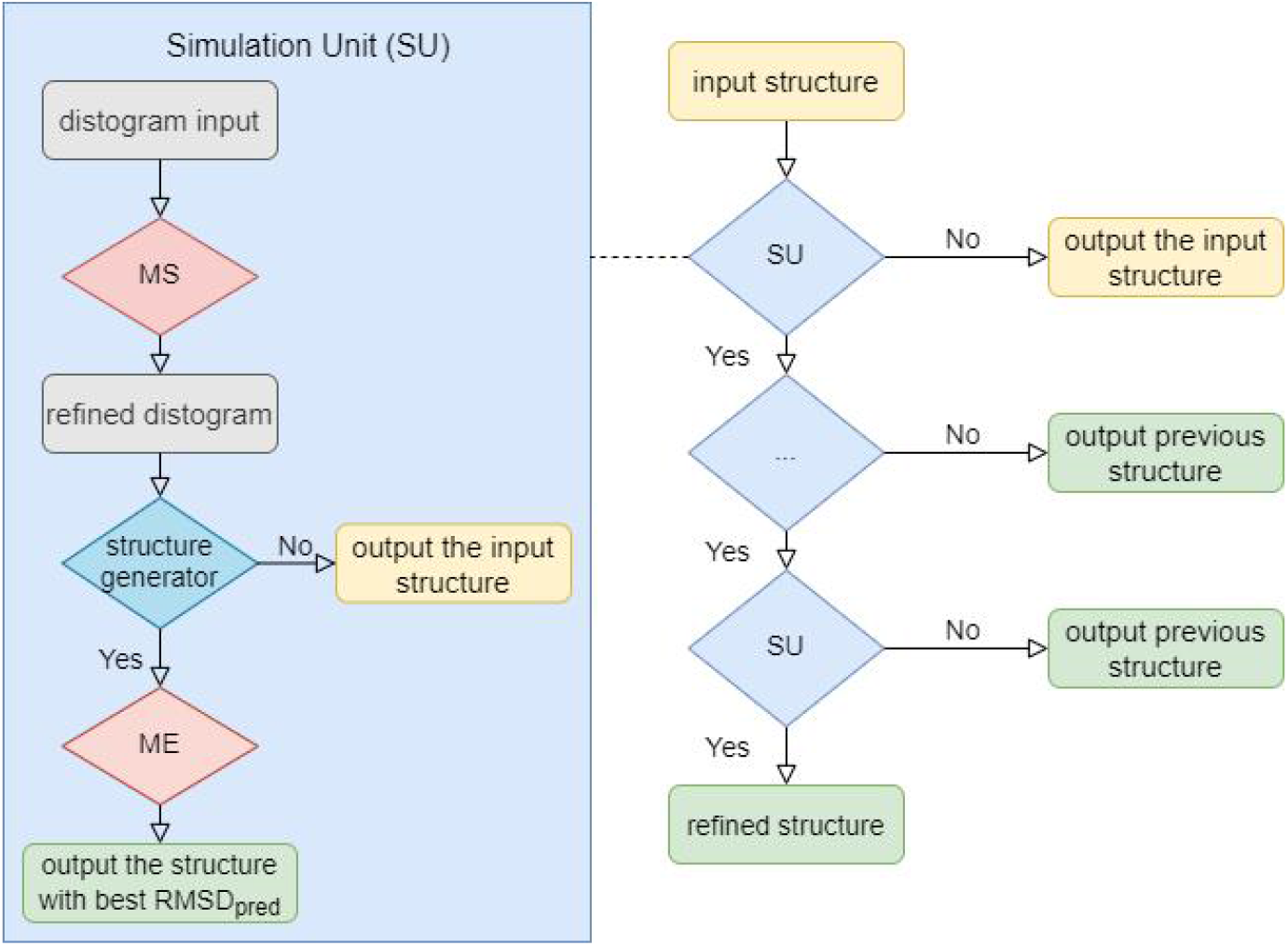
Pipeline of refinement. (right) and the simulation unit (SU) (left)

Considering the relatively small quantity of training data for each loop we used, this is acceptable. More loops and FOLD_M data should be introduced to the training set to solve this overfitting problem.

### Refinement using distogram

#### Model accuracy

We trained the model MS 34 epochs, and Figure 12 shows the performance of epoch 34 of MS using the test dataset. The left chart of Figure 12 is dRMSD_pred_ as a function of dRMSD_input_. The Pearson correlation coefficient of dRMSD_pred_ and dRMSD_input_ is 0.98, and the slope is 0.92, which is smaller than 1. Our model produces the distance matrix closer to the native structure compared to the input distance matrix. The right chart of Figure 12 is the distribution of ΔdRMSD, which is defined as dRMSD_input_ - dRMSD_pred_. 64% cases are getting improved with ΔdRMSD larger than 0, i.e., the accuracy of our MS model is 64%.

**Figure 12:**
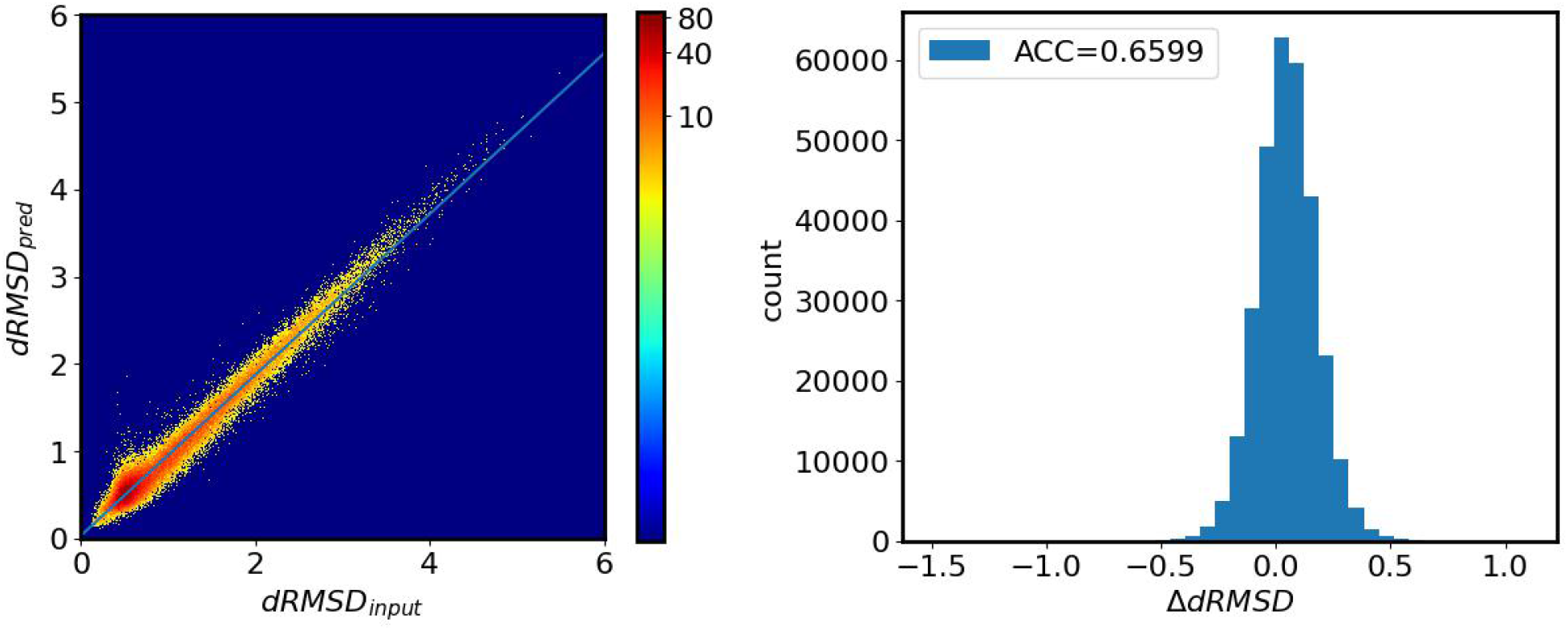
Performance of sampling model (MS) at epoch 34. Left chart: the 2D histogram of dRMSD_pred_ as a function of dRMSD_input_. The population is indicated as the color, and the scale is shown in the color bar. The linear regression result is shown as the blue line, with Pearson correlation coefficient as 0.98, slope as 0.92, and intercept as 0.04. Right chart: the distribution of ΔdRMSD, which is defined as dRMSD_input_ - dRMSD_pred_. 64% cases are getting improved with ΔdRMSD larger than 0, i.e., the accuracy of our MS model is 64%.

The best epochs are EP21 and EP13 for ME_resNet and ME_resNet_SD model based on the validation dataset. Figure 13 shows the performance of ME_resNet_EP21 and ME_resNet_SD_EP13 using the test dataset. The top two figures show the 2D histogram of predicted RMSD as a function of ground true RMSD. The Pearson correlation coefficients are 0.866 and 0.858, and the slopes are 0.775 and 0.780 for ME_resNet and ME_resNet_SD, respectively. We notice that both models are trending to predict the RMSD smaller than the ground true RMSD. The bottom chart of Figure 13 shows the difference of true RMSD and predicted RMSD, i.e., RMSD_true_-RMSD_pred_, and the ME_resNet_SD is more prospective to predict RMSD smaller than ground true RMSD. When using the energy model to refine the structure, the absolute RMSD is less meaningful than the relative RMSD. The key point is by given two structures whether the energy model could pick out the better one. We test our models by randomly picking 305019 pairs of fragments from the test dataset, and each pair contains two conformations of the same fragment. The accuracy is shown in Table 1. The models reach an accuracy of 97% if the difference of ground true RMSD is larger than 2 Å, and the accuracy drops to 94% for range [1.0 Å - 2.0 Å), 89%-90% for range [0.5 Å - 1.0 Å), and 70% for range smaller than 0.5 Å. Since the protein structure is flexible, it is difficult to distinguish two structures that are very close to each other.

**Figure 13:**
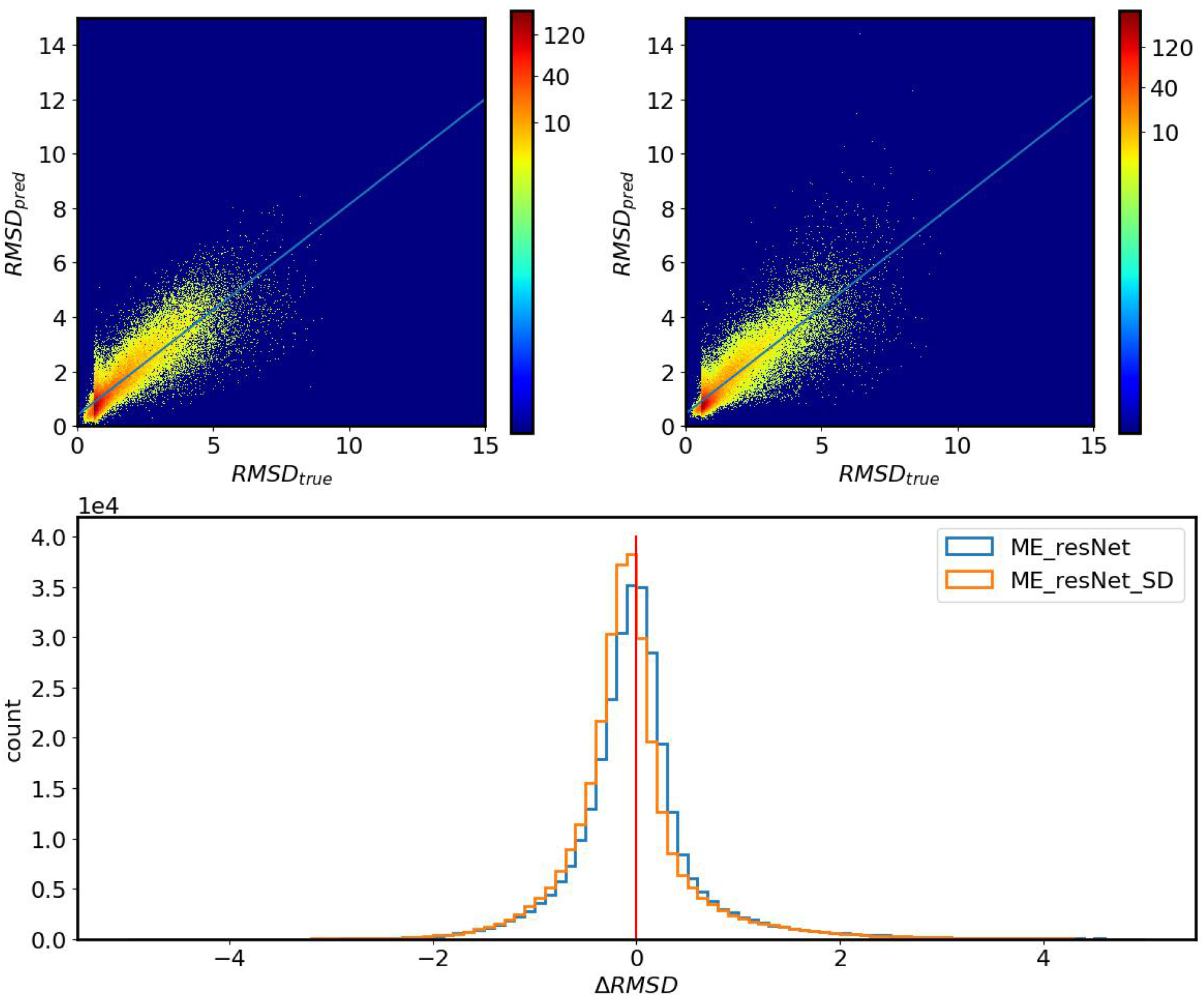
Performance of energy model. The top figures (left: ME_resNet_EP21, right: ME_resNet_EP13) show the 2D histogram of predicted RMSD as a function of ground true RMSD. The population is indicated as the color, and the scale is shown in the color bar. The linear regression result is shown as the blue line, with Pearson correlation coefficient as 0.866 and 0.858, slope as 0.775 and 0.780, and intercept as 0.389 and 0.444 for ME_resNet_EP21 and ME_resNet_EP13, respectively. The bottom Figure shows the difference of RMSD, i.e., ΔRMSD = RMSD_true_ -RMSD_predict_, the red line indicates 0.

**Table 1:** The accuracy of energy model ME_resNet_EP21 and ME_resNet_SD_EP13 to compare pairwise conformations of fragments. The data are randomly picked from the test dataset. The accuracy is defined as how many pairs that the sign of ΔRMSD_pair_ is the same as the sign of ΔRMSD_true_.

| $\Delta\text{RMSD}_{\text{pair}}$ (Å) | ME_resNet_EP21 | ME_resNet_SD_EP13 |
| --- | --- | --- |
| [0, 0.5) | 0.7029 | 0.6941 |
| [0.5, 1.0) | 0.8991 | 0.8885 |
| [1.0, 2.0) | 0.9405 | 0.937 |
| > 2.0 | 0.9762 | 0.9725 |

#### Simulation result

We run two sets of refinement experiments, both with MS_EP34 to generate refined distogram, but using different quality control energy models, one with ME_resNet_21and the other one with ME_resNet_SD_13. Table 2 shows the performance of these two experiments. The accuracy is defined whether the RMSD of the structures with top 1 or top 5 predicted RMSD is lower than the input structure from the 100 steps simulation. ME_resNet has better performance than ME_resNet_SD with top1 accuracy of 0.6194, i.e., 61.94% fragments get improved by using our model. And the top 5 accuracy reaches 0.7161 for both ME_resNet and ME_resNet_SD model. In the six-residues fragments, we define a well-structured fragment (SS) as at least four residues of the native structure are helix or sheet assigned by DSSP[40]. If more than two residues of the native structure are turn or coils, we define it as a non-structured fragment (nonSS). Our models have a decent performance on the well-structured fragments with top 1 (top 5) accuracy as 0.7125 (0.80) for ME_resNet and 0.7250 (0.80) for ME_resNet_SD. For non-structured fragments, the top 1 (top 5) accuracy is only 0.5135 (0.6216) for ME_resNet and 0.4459 (0.6216) for ME_resNet_SD, respectively. The ME_resNet has better top-1 accuracy on the non-structured fragments than ME_resNet_SD. We also calculate the success rate in the simulation experiments, which is defined as the average percentage of generated structures among the number of attempts, i.e., 100 steps for each fragment. The success rate is 89.22% for ME_resNet and 93.38% for ME_resNet_SD. The ME_resNet_SD has a better success rate, but lower accuracy shows that the ME_resNet_SD model learns more “nature-wise” energy function, which could pick a reasonable distogram to generate a structure. But the model is not well trained yet, which requires either longer training or better hyperparameters.

**Table 2:** The accuracy of simulation experiments by using different energy models. The accuracy is defined as that the RMSD of the structure with top1 or top5 predicted RMSD is lower than the input structure. SS accuracy shows the accuracy of fragments with more than four residues of the native structure are helix or sheet, and nonSS is the accuracy of fragments with more than four residues of the native structure are random coil. The secondary structure is assigned by DSSP.

| Accuracy | Top 1 | Top 5 | SS<sub>top1</sub> | SS<sub>top5</sub> | nonSS<sub>top1</sub> | nonSS<sub>top5</sub> |
| --- | --- | --- | --- | --- | --- | --- |
| ME_resNet | 0.6169 | 0.7142 | 0.7125 | 0.8000 | 0.5135 | 0.6216 |
| ME_resNet_SD | 0.5909 | 0.7143 | 0.7250 | 0.8000 | 0.4459 | 0.6216 |

Figure 14 shows the violin plot of Δ RMSD (RMSD_true_-RMSD_pred_) for initial structures, successively refined structures (RMSD smaller than initial structure), and the failed structures (RMSD larger than initial structure). For the initial structure, both models have a similar distribution, and ME_resNet_SD has more population around 0 Å but a lower mean value of Δ RMSD as -0.0321 Å compared to ME_resNet (−0.0070). For the successful cases, the distribution is mainly located around 0 but with a high outlier for ME_resNet; the distribution of ME_resNet_SD is bimodal with one peak located about 0 Å and the other peak around 1 Å. The bimodal distribution of ME_resNet_SD on the successful cases shows that this model has unstable performance on different pdb structures. For some structures, ME_resNet_SD could predict RMSD close with the true RMSD, which corresponds to the peak at 0 Å in the violin plot; but for some structures, the model trends to predict smaller RMSD than the true RMSD, which is shown in the peak at 1 Å in the violin plot. For these structures, the model ME_resNet_SD could give accuracy pairwise RMSD, i.e., between two conformations, the model could figure out the one with lower RMSD. In the failed cases, both models tend to predict smaller RMSD.

**Figure 14:**
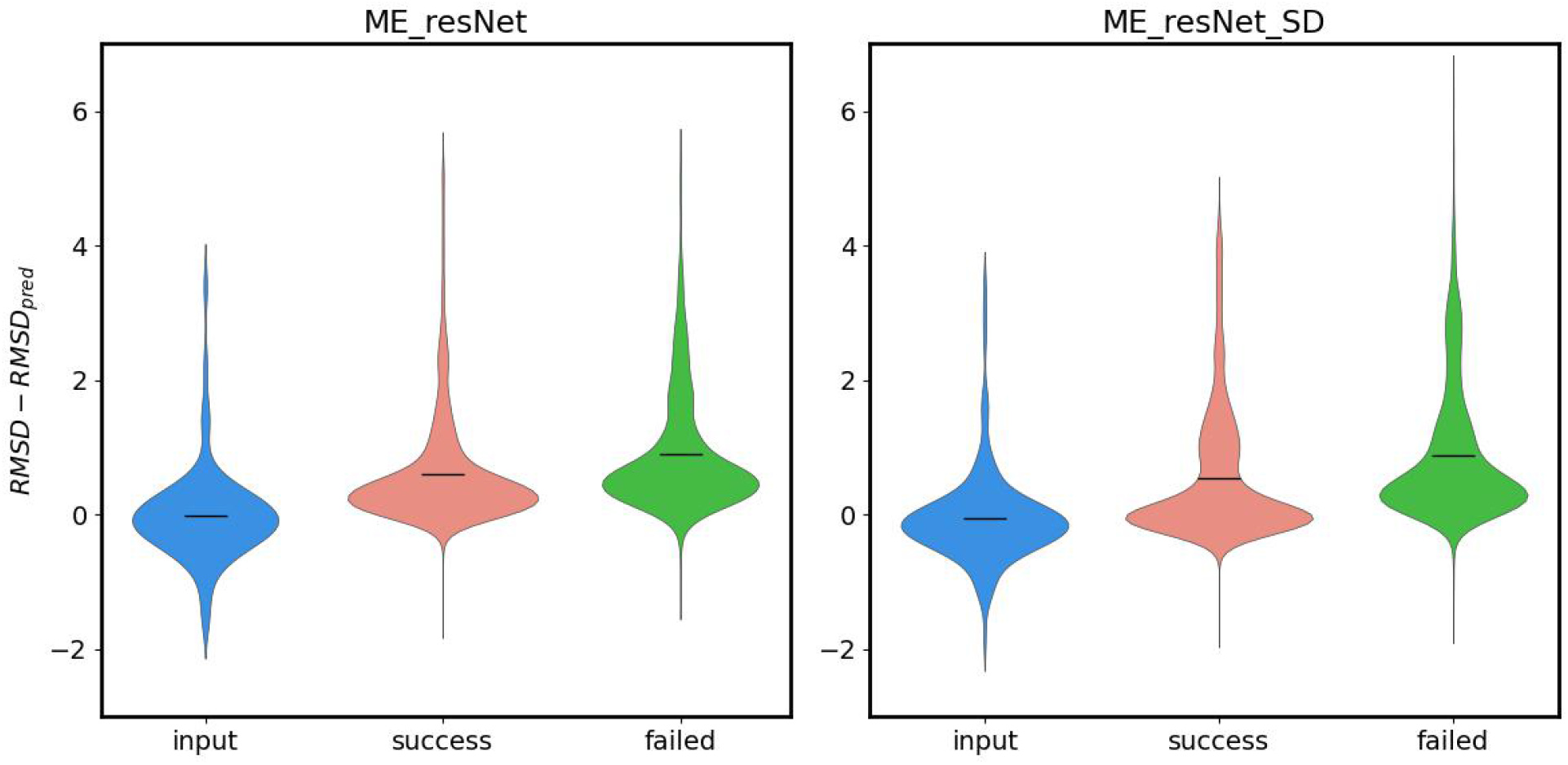
Violin plot of Δ RMSD of initial and refined structures of the 160 simulation experiments by using ME_resNet (left) and ME_resNet_SD (right) as the energy function. Blue violins show the Δ RMSD distribution of initial structures, the red ones show the result of successively refined structures (RMSD smaller than initial structure), and the green ones show the result of the structures with RMSD larger than the initial structure. The horizontal black lines show the mean value of the ΔRMSD, which are -0.0070, 0.6137, 0.8897, -0.0321, 0.5595, and 0.8613 from left to right.

Figure 15 shows one successful case (pdb ID: 1o6v, chain: A, fragment residue index: 330-336) from the 154 simulation experiments. In the first few steps, the structure gets improved towards the native structure, with RMSD decreasing dramatically for both models. Comparing to the initial structure with RMSD at 2.36 Å, both models output an improved structure with RMSD at 0.37 Å (ME_resNet) and 0.79 Å (ME_resNet_SD) with the lowest predicted RMSDs at 0.0039 Å (ME_resNet) and 0.47 Å (ME_resNet_SD). Among the 100 steps, the lowest RMSD is 0.26 Å (ME_resNet) and 0.40 Å (ME_resNet_SD). The model ME_resNet performs better in this case, especially after step 10, the model can generate fragments stable around 0.5 Å, but ME_resNet_SD has more fluctuation with the RMSD of the predicted structures from 0.40 Å to 2.00 Å. The main problem in this experiment is that both sampling and energy models are trained using the structures sampled from torsion sampling models. The models cannot handle the structures generated using the refined distogram, resulting in an unstable performance after the first few simulation experiments. To solve this problem, we plan to generate more structures by using our distogram algorithm and mixing these structures with the previous training data to retrain the models.

**Figure 15:**
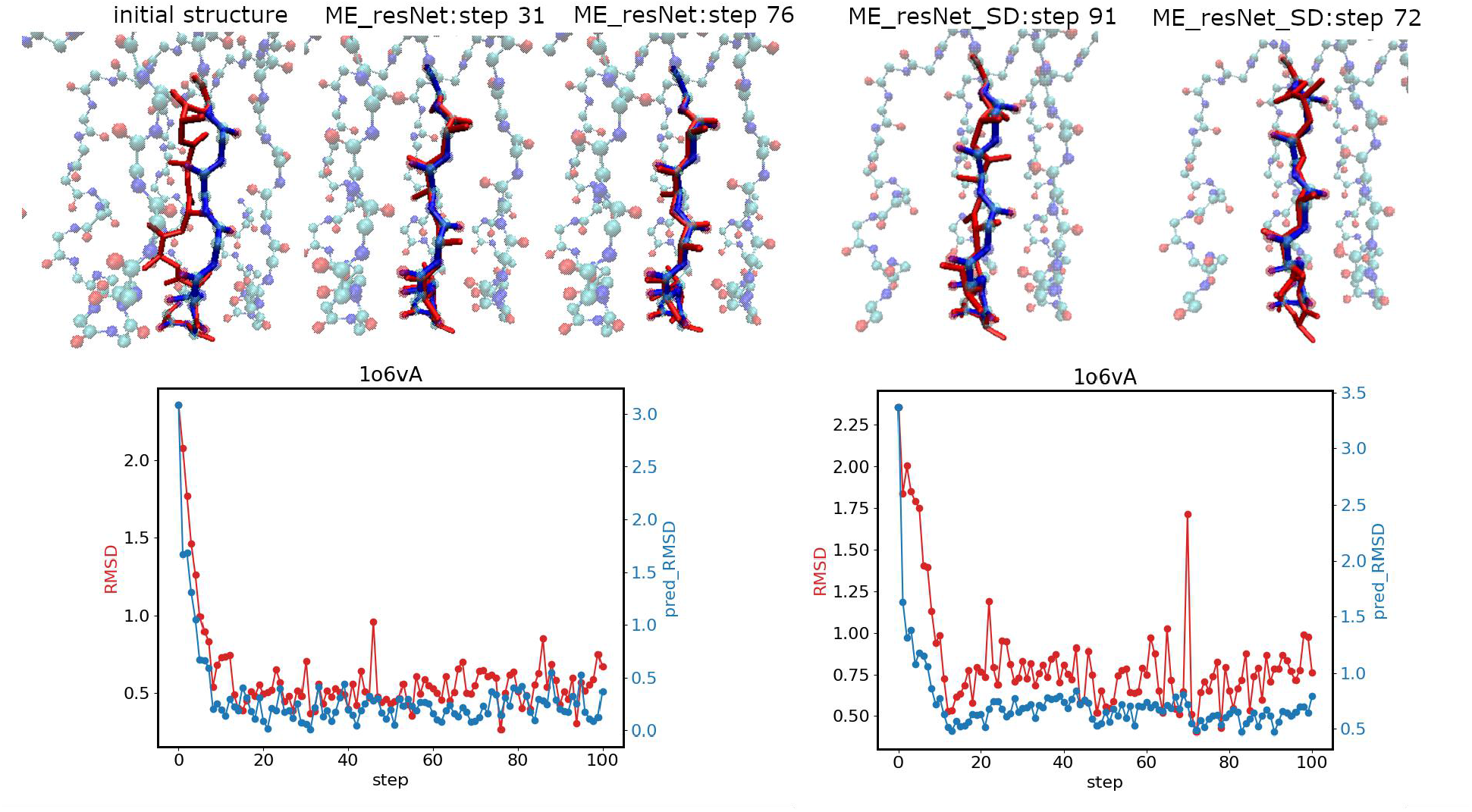
One successful case. (pdb ID: 1o6v, chain: A, fragment residues index: 330-336) from the 154 simulation experiments by using ME_resNet (left) and ME_resNet_SD (right) as energy function. The top row shows the snapshot of the initial/refined structures of the target fragment (red) compared with the native structure (blue). From left to right are initial structure, the structure (generated in step 31^_st_^ of 100 steps by using MS + ME_resNet) with best-predicted RMSD by ME_resNet; the structure (step 76^_th_^ of 100 steps by using MS + ME_resNet) with best true RMSD; the structure (generated in step 91^_st_^ of 100 steps by using MS + ME_resNet_SD) with best predicted RMSD by ME_resNet_SD, and the structure (step 72^_nd_^ of 100 steps by using MS + ME_resNet_SD) with best true RMSD. The bottom charts show the true RMSD (red) and predicted RMSD (blue) of the structures in each step in the experiments MS+ME_resNet (left) and MS+ME_resNet_SD (right).

## Conclusions

With well-designed deep learning architecture, our torsion sampling and energy scoring models show good prediction accuracy. The torsion sampling model performs a good balance of efficiency and diversity, while the energy scoring model gives a high linear correlation between predicted and real energy scores. Asymmetric loss function was found to optimize the model better than the MSE loss function. In the folding simulations with ten loops from the D3 dataset, the resultant average RMS is 1.39Å, which is a state-of-art result for loops with 16 residues. Although the performance cannot be generalized well to test loops out of the training set, this methodology shows good potential to solve the loop modeling problem considering the small quantity of training data we used. A more extensive training dataset and proper techniques to help generalization should improve the models. Besides, the distogram-guided refinement method gives good results of refining fragment to lower RMS scores. Our distogram-guided refinement procedure improved 61.69% out of 154 fragments in our distogram refinement test dataset. In the example we showed, the loop with an initial RMS of 2.36Å was refined to only 0.26Å. The current models in the distogram-guided refinement were trained for general fragments, and performance is not standout for loop modeling. We will use a dataset contains only loop structures to re-train the model for loop modeling.

## Acknowledgments

Research reported in this publication was supported by the National Institute of General Medical Sciences of the National Institute of Health under award number R01GM126558. The content is solely the responsibility of the authors and does not necessarily represent the official views of the National Institutes of Health.

